# *Chlamydia trachomatis* Encodes a Dynamic, Ring-Forming Bactofilin Critical for Maintaining Cell Size and Shape

**DOI:** 10.1101/2020.10.19.346502

**Authors:** Mary R. Brockett, Junghoon Lee, John V. Cox, George W. Liechti, Scot P. Ouellette

## Abstract

Bactofilins are polymer-forming cytoskeletal proteins that are widely conserved in bacteria. Members of this protein family have diverse functional roles such as orienting subcellular molecular processes, establishing cell polarity, and aiding in cell shape maintenance. *Chlamydia* species are obligate intracellular bacteria that undergo a developmental cycle alternating between an infectious, non-dividing EB and a non-infectious, dividing RB. As *Chlamydia* divides by a polarized division process, we hypothesized that BacA_CT_ may function to establish polarity in these unique bacteria. Using sequence alignment to the conserved bactofilin domain, we identified a bactofilin ortholog, BacA_CT_, in the obligate intracellular pathogen *Chlamydia trachomatis*. Utilizing a combination of fusion constructs and high-resolution fluorescence microscopy, we determined that BacA_CT_ forms a dynamic, membrane-associated, ring-like structure in *Chlamydia’s* replicative RB form. Contrary to our hypothesis, this filamentous ring structure is distinct from the microbe’s cell division machinery and does not colocalize with septal peptidoglycan or MreB, the major organizer of the bacterium’s division complex. Bacterial two-hybrid assays demonstrated BacA_CT_ interacts homotypically but does not directly interact with proteins involved in cell division or peptidoglycan biosynthesis. To investigate the function of BacA_CT_ in chlamydial development, we constructed a conditional knockdown strain using a newly developed CRISPR interference system. We observed that reducing *bacA_CT_* expression significantly impacted chlamydial cell size and morphology. Normal RB morphology was restored when an additional copy of BacA_CT_ was expressed *in trans* during knockdown. These data reveal a novel function for chlamydial bactofilin in maintaining cell shape in this obligate intracellular bacterium.

**IMPORTANCE:** *Chlamydia* is an ancient, obligate intracellular bacterium with a unique biphasic developmental cycle. As a result of its evolution within the osmotically stable environment of a host cell, *Chlamydia* has lost its dependence on side-wall peptidoglycan, and maintains only a fraction of the components thought to be required for regulating bacterial cell size and division. As such, very little is known about how *Chlamydia* species carry out these critical processes in the absence of a stabilizing peptidoglycan layer. In the current study, we identify a novel cytoskeletal element, termed a bactofilin, that is critical for maintaining the morphology of the bacteria. Using state-of-the-art genetic techniques for this organism, we demonstrate that chlamydial bactofilin forms a dynamic ring structure independent of the microbe’s division machinery and that abrogating its expression level using CRISPR interference results in abnormal morphologic forms. These findings enhance our understanding of chlamydial biology and bactofilins more generally.

## INTRODUCTION

*Chlamydia trachomatis* is the causative agent of non-congenital infectious blindness (trachoma biovar: serovars A-C), the leading cause of bacterial sexually transmitted genital tract infections (genital tract biovar: serovars D-K), and the cause of invasive urogenital or anorectal infection (lymphogranuloma venereum biovar: serovars L1–L3) (1). Untreated genital infections can result in urethritis in men and cervicitis, pelvic inflammatory disease (PID), and ectopic pregnancy in women. The vast majority of this disease burden in the US is borne by young women between the ages of 15-24 (2). *C. trachomatis* is an obligate intracellular bacterium that has a unique biphasic developmental cycle (1). The infectious, non-replicative elementary body (EB) attaches to and is endocytosed by susceptible host cells. Within a host-derived, membrane bound compartment (inclusion), the EB differentiates into its replicative form, the reticulate body (RB). The RB grows and divides within the inclusion before asynchronously differentiating into EBs to exit the host cell.

Most bacterial species utilize the glycopolymer peptidoglycan to maintain cell shape and membrane integrity(3). The peptidoglycan layer is highly dynamic and is actively shaped throughout bacterial growth and division. Localized peptidoglycan synthesis and hydrolysis is directed by filament-forming, cytoplasmic cytoskeletal proteins (4). The actin-like protein MreB directs peptidoglycan synthesis around the cell periphery while the tubulin-like protein FtsZ initiates peptidoglycan synthesis at the site of division (5). FtsZ polymers form a ring that provides the mechanical force for constriction of the cell and recruits other key Fts family proteins to the site of division.

As a result of more than 700 million years of adaptation to an intracellular environment, *Chlamydia* species are physiologically unique in that they lack a number of genes that are essential in other bacteria, such as *ftsZ* (6, 7). It was recently determined that *Chlamydia* utilizes MreB to direct septal peptidoglycan biosynthesis and that MreB function is essential for *Chlamydia* division (8, 9). Paradoxically, *C. trachomatis* does not generate side-wall peptidoglycan and maintains only a thin septal band of peptidoglycan at its division septum (9, 10). As peptidoglycan acts as a major structural support between the inner and outer membranes in Gram-negative bacteria, it is presently unclear how *Chlamydia* species maintain membrane integrity and provide structural support for transmembrane complexes in their replicative forms.

In addition to FtsZ and MreB, a new bacterial protein family of self-assembling, polymer-forming landmark proteins has been described; bactofilins. These intermediate filament-like cytoskeletal proteins have been identified in several bacterial species (11–14) and perform a number of critical cellular functions such as establishing cell polarity (14), maintaining cell shape by directing localized peptidoglycan biosynthesis and degradation enzymes (15), and aiding in chromosomal segregation during cell division (16). In this study, we identified and characterized a chlamydial bactofilin ortholog, BacA_CT_. Using a combination of fixed and live-cell imaging in conjunction with a recently described CRISPRi technique, we demonstrate that BacA_CT_ functions in maintaining membrane integrity outside of the chlamydial septum by forming dynamic, ring-like structures whose dimensions are similar to the previously characterized chlamydial peptidoglycan ring.

## Results

### The chlamydial gene, *ct276* /*ctl_0528*, encodes a bactofilin ortholog

Recently, a number of studies have revealed the importance of intermediate filament-like proteins in bacteria. This class of proteins has been designated bactofilins, and they have been associated with diverse functions such as determining cellular polarity and maintaining cell shape (14). Given that pathogenic *Chlamydia* species divide by a polarized division mechanism (17–19), we hypothesized that they encode a bactofilin homolog that could serve to establish polarity in this unique organism. Bactofilins retain a conserved domain, DUF583 (20), that encodes for a β-helical core domain essential for the polymerization of protein monomers (11–14). We used this domain to bioinformatically search for a chlamydial homolog, and our analysis revealed that the gene *ct276*/ *ctl_0528* encodes a 194 amino acid protein with this conserved domain. Homologs of this gene are present in all pathogenic species of *Chlamydia* examined and are encoded 3’ to the *dnaA2* gene (**Fig. 1A, Fig. S1**). Interestingly, no homolog for BacA_CT_ is encoded in the genomes of any of the ‘Chlamydia-related’ bacteria that make up the other members of the order Chlamydiales, such as *Parachlamydia acanthamoebae*, *Simkaniae negevensis*, and *Waddlia chondrophila* Pathogenic *Chlamydia* species differ from these *Chlamydia*-related bacteria in one key way: they lack a peptidoglycan sacculus.

**Figure 1.**
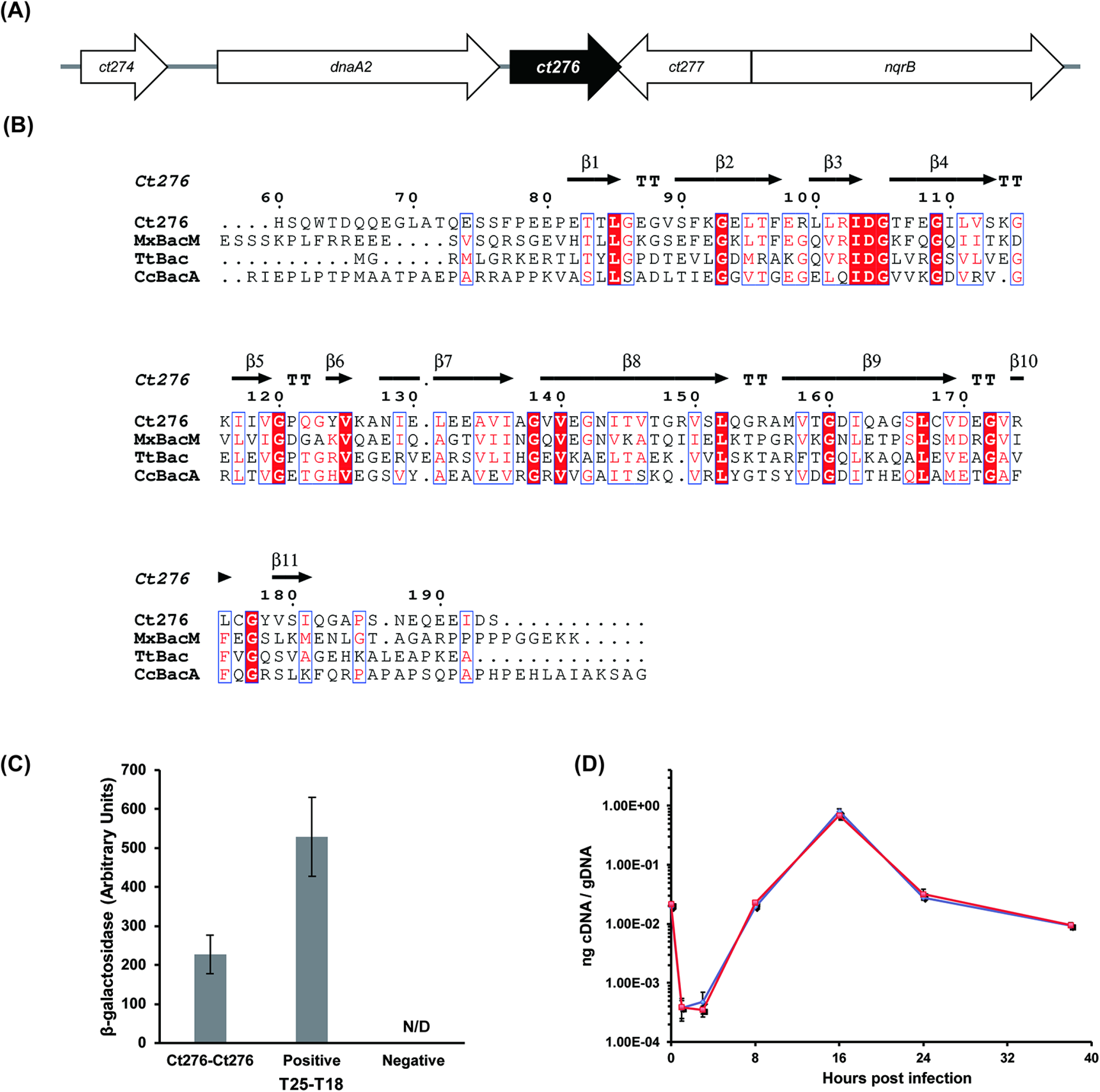
The chlamydial gene CT276 / CTL_0528 (*bacA_CT_*), encodes a bactofilin protein. **(A)** Illustration of the *bacA_CT_* locus from *C. trachomatis* serovar D UW-3/Cx. Gene order and orientation is highly conserved in all pathogenic *Chlamydia* species examined. **(B)** Amino acid alignments of *bacA_CT_* with other DUF583 domain containing bactofilins (*Myxococcus xanthus* BacM, and *Thermus thermophilus* Bac, *and Caulobactor crescentus* BacA). The DUF583 domain, a characteristic structural region of all bactofilin proteins, is composed of sequential beta sheets (black arrows). **(C)** The capacity for BacA_CT_ monomers to directly interact which each other was quantified utilizing a bacterial adenylate cyclase-based two hybrid (BACTH) system. Readouts were assessed by β-galactosidase assay. A positive control is an interaction between T25-Zip and T18-Zip, the GCN4 leucine zipper motif, and a negative control is the lack of interaction between T25 and T18-Ct276. The entire assay was conducted in biological triplicate and error bars represent standard deviations of the mean. N/D, Not detected. **(D)** Expression profile of *bacA_CT_* during the developmental cycle of *C. trachomatis*. Each data point represents the mean of three technical replicates, and data are shown from two independent biological replicates (red vs black).

Sequence alignments and *in silico* protein prediction models indicate that the β-helical core is highly conserved at the amino acid (**Fig. 1B**, >57% identity/similarity) and structural level **(Fig. S1)**. As the closest structural homolog with a solved crystal structure in the Protein Data Base (PDB) is the bactofilin encoded by *Caulobactor vibrioides* (*bacA*), we chose to rename the gene encoding the chlamydial bactofilin as *bacA_CT_*. Since the capacity to form filaments is a characteristic of all bactofilin proteins, we tested the ability of BacA_CT_ to interact homotypically using the bacterial adenylate cyclase-based two hybrid (BACTH) system. We found that BacA_CT_ interacts with itself **(Fig. 1C)**, as expected, similar to results previously reported in a much larger protein-protein interaction study that utilized a yeast two hybrid system (21). Based on these data, we conclude that BacA_CT_ is a canonical bactofilin.

Given that *Chlamydia* is characterized by its unique developmental cycle, we monitored the transcription of *bacA_CT_* to determine whether its expression pattern was associated with early cycle events (i.e. EB to RB differentiation), mid cycle events (i.e. growth and division of the RB), or late cycle events (i.e. RB to EB differentiation). We determined that *bacA_CT_* transcription peaks at 16h post-infection **(Fig. 1D)**, consistent with previous studies (22) and indicating that the protein may function in RB growth and division.

### BacA_CT_-mCherry polymers are dynamic, alternating between puncta and ring-like structures

Bactofilin proteins exhibit different functions in different bacterial species that can often be inferred based on their localization within a microbe. For example, BacA from *C. crescentus* localizes to a single pole and directs the cell division machinery within the bacterium (14), whereas CcmA in *Helicobacter pylori* localizes to a single face of the bacterium’s cell wall and directs peptidoglycan remodeling enzymes responsible for maintaining the microbe’s unique helical shape (23). We wanted to determine where BacA_CT_ localizes within *Chlamydia* RBs. To investigate the subcellular localization of BacA_CT_, we constructed C-terminal hexahistidine (6xH) and mCherry BacA_CT_ fusion constructs for expression in the *C. trachomatis* strain L2 434/Bu. Constructs were cloned into shuttle plasmids under the control of an anhydrous tetracycline inducible (aTc) promoter and transformed into *C. trachomatis*, as described previously (17, 24). Cell division genes are precisely regulated in bacteria, and their overexpression often causes fitness defects, including in *Chlamydia* (25, 26). To eliminate this possibility, we first examined whether the induction of BacA_CT_ fusion constructs affect the fitness of *C. trachomatis* transformants by quantifying the ability of the organism to complete its developmental cycle and generate infectious elementary bodies (or inclusion forming units, IFU). We found that the induction of our BacA_CT_ constructs resulted in no measurable differences in the number of IFU recovered at 44 hpi, regardless of the concentration of aTc **(Fig. S2)**, indicating that the induction of BacA_CT_ does not adversely affect progression through the developmental cycle.

We next utilized our BacA_CT_-mCherry expressing strain to observe filament formation and localization in actively growing chlamydial RBs via live-cell imaging microscopy. We observed that the fusion protein predominantly accumulated in puncta, filaments, or partial ‘ring-like’ structures **(Fig. 2A, B)**. A quantitative analysis of the spatial dimensions and fluorescence intensity associated with individual bactofilin structures in *Chlamydia* indicated that, when accounting for differences in volume, puncta forms displayed much higher mean fluorescence intensity measurements than filaments or ring-like structures **(Fig. 2C, Fig. S3, Video S1-3)**. We hypothesized that this might indicate that puncta may differ from filamentous / ring-like forms by the density of their monomeric bactofilin proteins. As these strains retain their parental BacA_CT_, we could not exclude the possibility that our fusion constructs simply tend to associate better with puncta than with rings / filamentous forms. However, an alternative explanation is that rather than two completely different structures, these forms are actually transient and interchangeable. To test this possibility, we conducted a series of time course studies to determine the dynamics of these structures in living cells. We observed that larger BacA_CT_ puncta are capable of transitioning into ring-like forms in the span of approximately 30 minutes **(Fig. 2D, Video S4)**, suggesting that, similar to the peptidoglycan ring-like structures inherent to pathogenic *Chlamydia* species (9), these structures are dynamic. Based on these data, we hypothesize that BacA_CT_ functions in stabilizing cell morphology during the division process.

**Figure 2:**
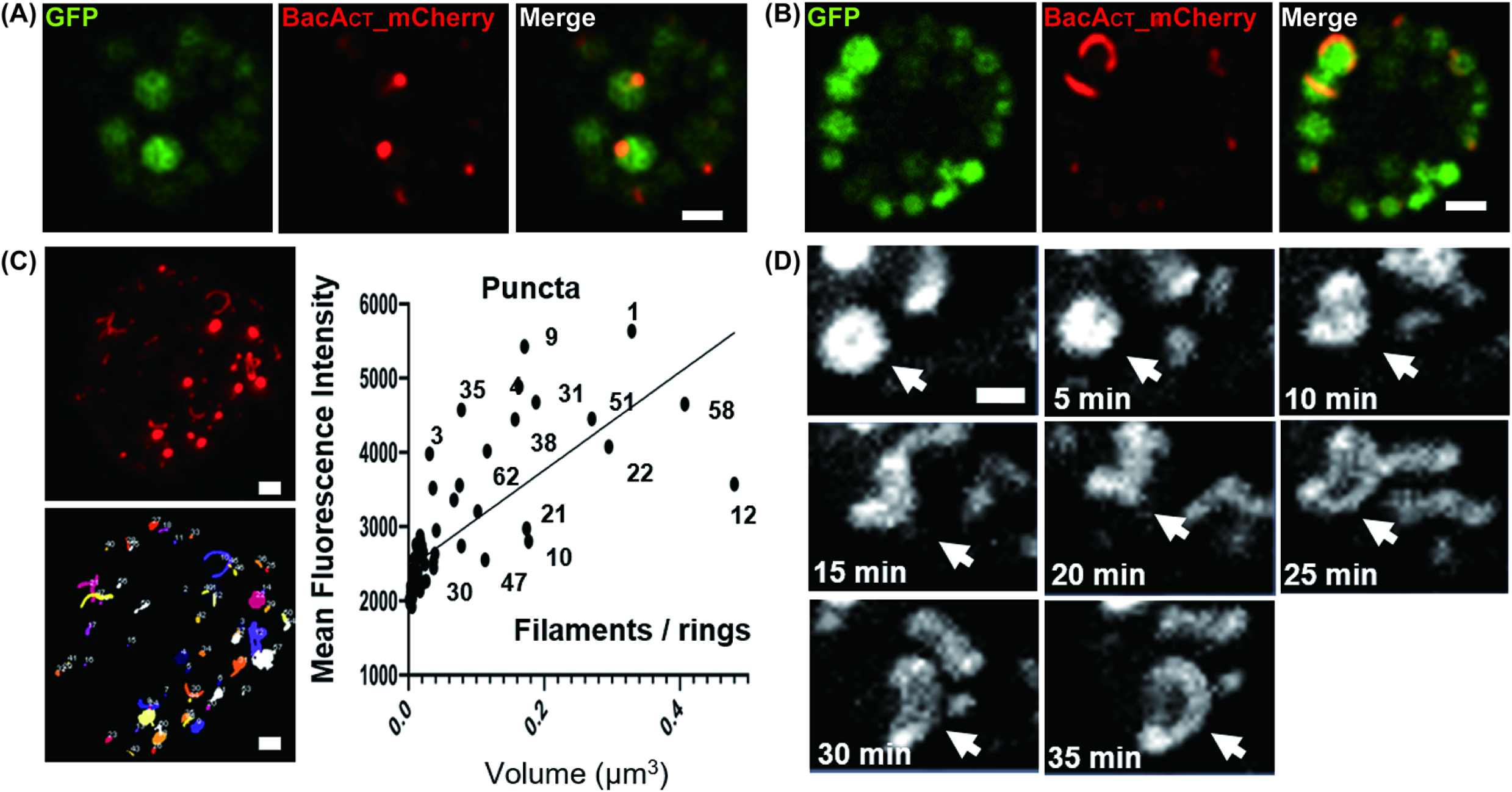
BacA_CT_ –mCherry localization is dynamic and alters between puncta, filaments, and ring-like structures. Live-cell imaging of *C. trachomatis* L2 434/Bu expressing a BacA_CT_ C-terminal mCherry fusion protein. Examples of polar puncta **(A)** and filaments and partial ring-like forms **(B)** are shown. Constitutively expressed cytoplasmic GFP is shown in green. Figures represent single imaging planes and are representative of ~30 fields of view observed. Scale bar, 1 μm. **(C)**Analysis of BacA_CT_ filament dimensions and density. Image is representative of 1 of 10 inclusions for which the mCherry positive objects were defined and mean fluorescence pixel intensities plotted against object volume. Scale bar, 1 μm. **(D)** Time-lapse microscopy of an expanding BacA_CT_ ring. Z-stacks were taken once every five minutes for thirty minutes and images presented are maximum intensity projections. The mCherry positive BacA_CT_ structure being tracked throughout the time course is designated by the white arrow. Scale bar, 0.5 μm.

### BacA_CT_ does not associate with cell division components

*Chlamydia* divides by a polarized division process. Given the dynamic nature of the BacA_CT_ rings and their apparent similarity in characteristics to the PG ring, we hypothesized that BacA_CT_ may function in the establishment of polarity and thus interact/intersect with the division machinery. We began by examining *C. trachomatis* BacA_CT_ localization during the initial stages of the developmental cycle. HeLa cells were infected with the transformant, and expression of the 6xH construct was induced at 6 hpi.

Infected cells were fixed to preserve their polarity at 10.5 hpi, a time when *C. trachomatis* is in its initial division (18), and processed for immunofluorescence analysis. We observed that full-length BacA_CT__6xH accumulated at or near the membrane of chlamydial RBs in short filamentous structures **(Fig. 3A)**. Filaments were not restricted to a particular site within the RB, exhibiting a roughly equal distribution between the budding daughter cell (upper panel – 29%), the opposite pole of the dividing RB (middle panel – 31%), and in association with the division plane (lower panel −40%). Thus, BacA_CT_ forms membrane-associated filaments that do not appear to preferentially localize to any particularly site in the dividing RB. However, we cannot exclude that there is a preference for an initial assembly site based on uncharacterized context-dependent cues.

**Figure 3.**
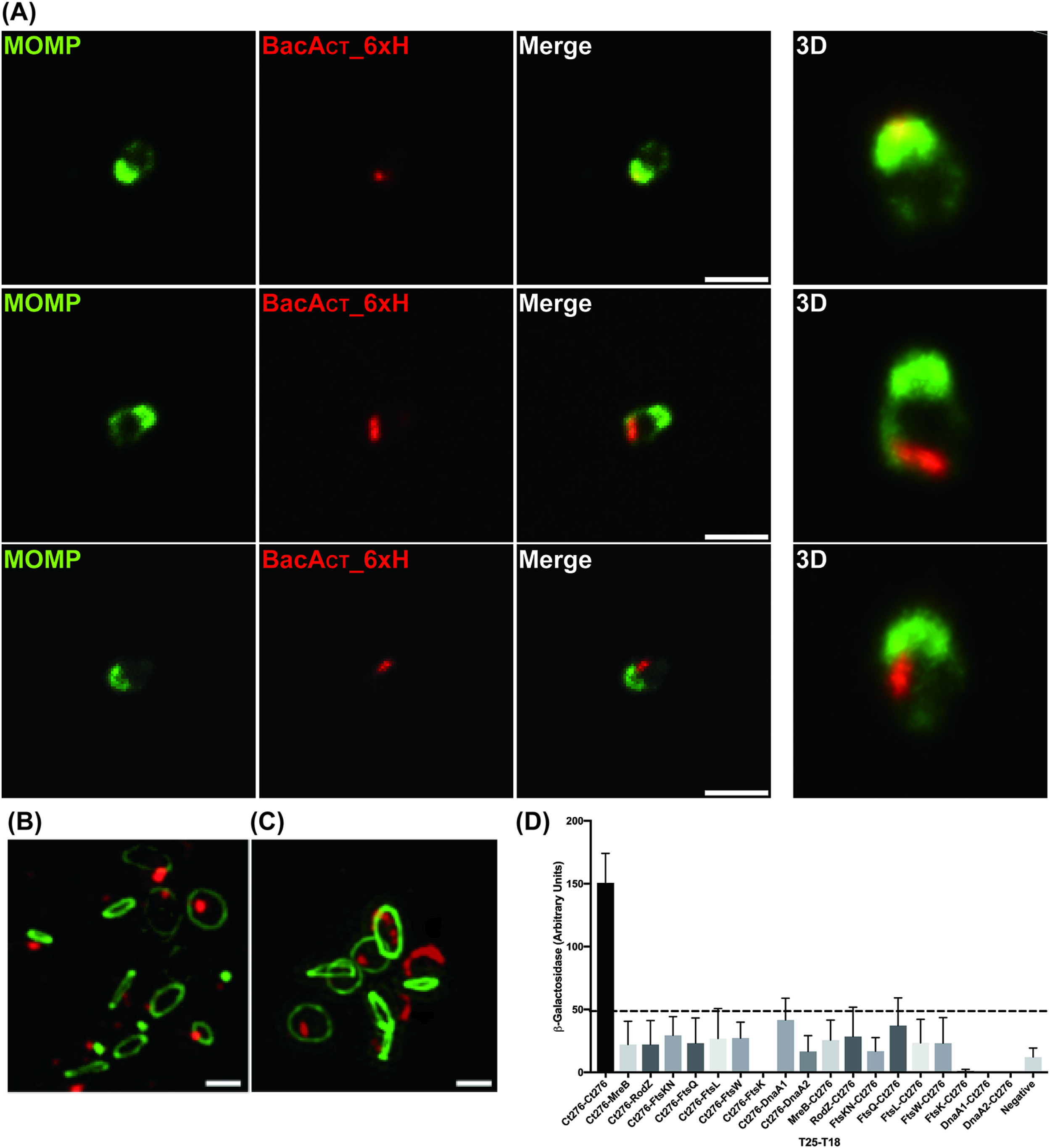
BacA_CT_ forms filaments that do not colocalize or interact with peptidoglycan or proteins associated with cell division in *Chlamydia* species. **(A)***C. trachomatis* strain L2 434/Bu 10.5 hours post-infection expressing BacA_CT_ with a six-histidine (6xH) tag at the C-terminus. RBs shown are undergoing the first step of their division process: polar expansion. The images were acquired on a Nikon Ti2 spinning disc confocal microscope using a 60X objective. Single imaging planes and maximum intensity projections are shown. Images are representative of over 50 imaging fields. Scale bar, 0.5 μm. **(B-C)** Super resolution microscopy of *C. trachomatis* expressing BacA_CT_-6xH (red) 18 hours post infection. Cells were co-incubated with the PG-intercalating agent EDA-DA (1 mM) and labeled with azide-conjugated Alexa Fluor 488 (green). Images are max intensity projections and representative of over 100 imaged fields. Scale bar, 1 μm. **(D)** BACTH and β-galactosidase assays were performed for BacA_CT_ interactions with proteins related to cell division and PG synthesis. The experiments were performed as described in Materials and Methods. The negative control was the lack of interaction between T25 and a mixture of T18-MreB and T18-FtsKN. The dashed line represents the cutoff for positive interactions as determined by five times the negative control levels. Data represent three independent biological replicates, and bars represent standard deviations from the mean.

We next evaluated BacA_CT_ localization with respect to the forming division septum, as indicated by the chlamydial PG structure, during the middle stages of the developmental cycle. We utilized the PG-labeling, D-amino acid dipeptide probe EDA-DA (9, 10), which is incorporated into newly synthesized bacterial PG by UDP-N-acetylmuramoyl-tripeptide-D-alanyl-D-alanine ligase (MurF) (9, 10, 27). An examination of the localization of BacA_CT_ filaments in RBs within fixed, *Chlamydia*-infected cells at 18 hpi revealed the presence of both filaments as well as punctate structures, similar to our observations at the early stage of the developmental cycle and to our live-cell imaging data (**Fig. 2)**. However, neither filaments nor puncta appeared to consistently co-localize with PG rings **(Fig 3B-C)**. As further evidence that BacA_CT_ does not directly associate with the division apparatus, we also labeled MreB, a major organizer of the chlamydial division and PG biosynthesis machinery (8, 9, 17, 28). We have previously shown that the PG ring and MreB co-localize in *C. trachomatis* (9), and, in line with our PG results in Fig. 3, we similarly found no co-localization of BacA_CT_ with MreB_CT_ **(Fig. S3)**.

Finally, we utilized a bacterial adenylate cyclase-based two hybrid (BACTH) system to test BacA_CT_ interaction with several key components of the *C. trachomatis* divisome (MreB (8, 9, 17, 28), RodZ (29, 30), FtsK (8), FtsQ (26), FtsL (26), and FtsW) as well as with DnaA2 (encoded in an apparent operon with *bacA_CT_*), and DnaA1. We were unable to detect any interaction between BacA_CT_ and any of these proteins **(Fig. 3D)**, indicating that BacA_CT_ does not appear to influence cell division through direct protein-protein interactions. Overall, our data indicate that BacA_CT_ does not function directly in cell division.

### Inducible knockdown of BacA_CT_ alters chlamydial morphology

To further investigate the function of BacA_CT_, we down-regulated its expression using CRISPRi technology we recently developed for *Chlamydia* (31). A chlamydial expression plasmid was constructed encoding an aTc-inducible dCas9 ortholog from *S. aureus* (32) and a constitutively expressed gRNA targeting the 5’ region of *bacA_CT_*. This vector was transformed into *C. trachomatis* L2 and subsequently used for experimentation. As a control, we also created a transformant carrying a plasmid encoding only the aTc-inducible dCas9.

Cells were infected with the transformants, and nucleic acids were collected under uninduced and induced conditions at 11.5, 16.5, and 24 hpi. Transcript levels were measured for *bacA_CT_*, the co-transcribed *dnaA2*, as well as an unrelated gene, *euo*, expressed early in the developmental cycle and associated with stress responses in *Chlamydia* (22, 33, 34). Transcripts for *bacA_CT_* were significantly reduced after dCas9 expression was induced in the transformant carrying the gRNA targeting *bacA_CT_*, whereas no consistent biologically meaningful change (>2x) across time points was observed in transcript levels of the co-transcribed *dnaA2* or the unrelated *euo* **(Fig. 4A-C)**. Conversely, the transcript levels of *bacA_CT_* were decreased at each time point we assessed. Interestingly, we noted distinct morphological changes in organisms in which *bacA_CT_* was knocked down **(Fig. 4A-C)**. Upon knocking down *bacA_CT_* expression, *Chlamydia* RBs were enlarged and displayed irregular localization of the chlamydial Major Outer Membrane Protein (MOMP). In the dCas9 only control, no changes were observed in transcript levels for any of the measured genes, and we did not observe any morphological defects **(Fig. S4A, C)**. We also assessed effects of *bacA_CT_* knockdown on peptidoglycan labeling. Peptidoglycan localization appeared to be largely unaffected, although larger ring structures were noticeably coincident with the presence of aberrantly enlarged reticulate bodies that exhibited a budding morphology **(Fig. 4D, E; Fig. S5)**.

**Figure 4.**
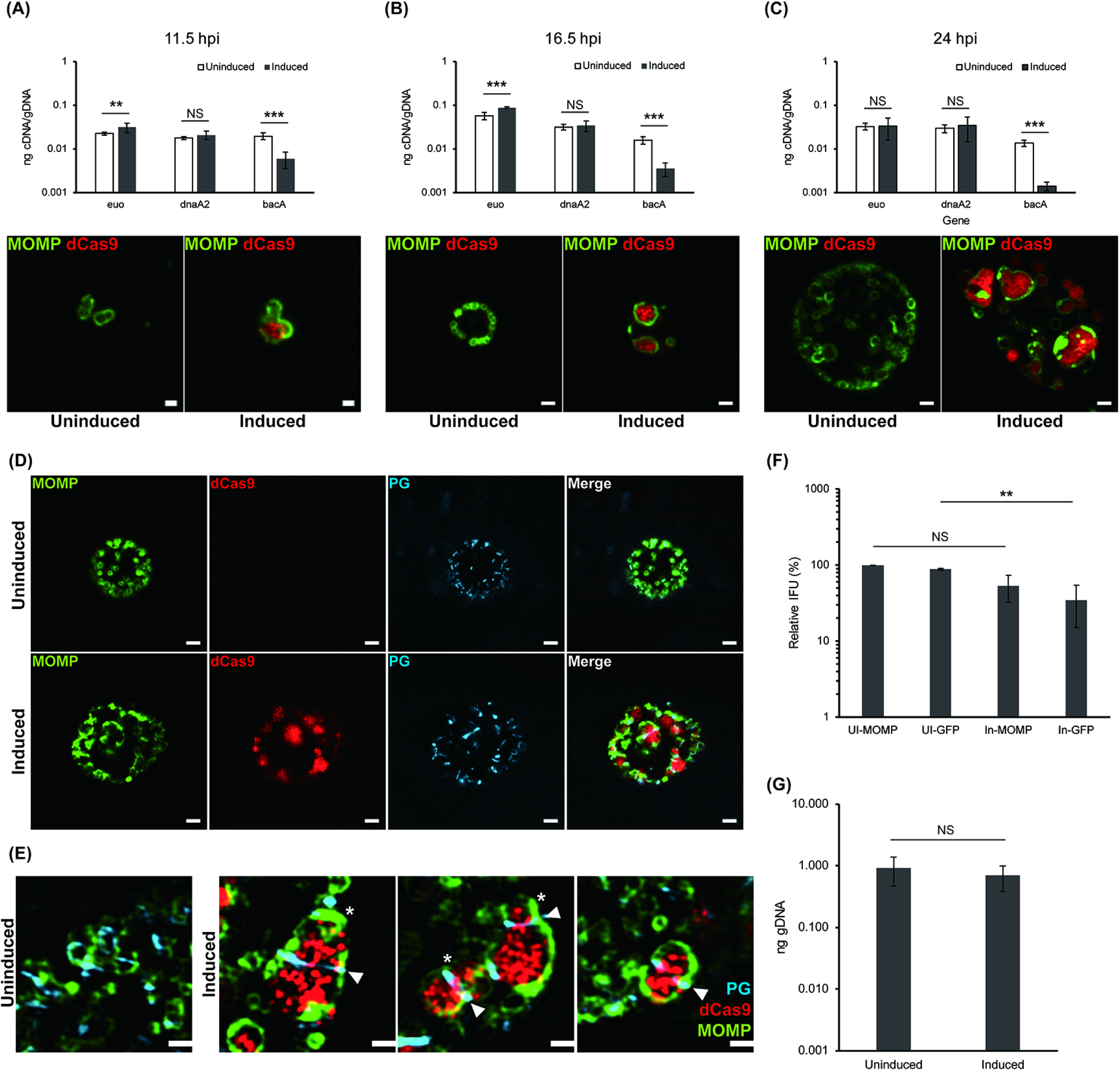
Conditional knockdown of *bacA_CT_* expression alters the morphology of *C. trachomatis*. *C. trachomatis* without plasmid (-pL2) was transformed with the aTc-inducible vector expressing a guide RNA targeting the 5’ region of *bacA_CT_* and the mutated *Staphylococcus aureus* Cas9, which cannot cleave DNA (dCas9) (31). After induction, transcript levels of *bacA* were measured by RT-qPCR at 11.5, 16.5, and 24 hpi **(A-C)**. The RT-qPCR was performed as described in Materials and Methods. In addition, immunofluorescent assay (IFA) was performed to check the induction of dCas9 and the morphology of the bacteria as assessed by labeling of the major outer membrane protein (MOMP) **(A-C)**. The merged channel is shown for each treatment. The PG was also labeled in live cells as described in Materials and Methods, and we performed fixation and labeling with anti-dCas9 antibodies for IFA **(D, E)**. For (D) and (E), arrowheads indicate the PG septum whereas * indicate budding daughter cells. Images were acquired on a Zeiss Imager.Z2 equipped with an Apotome2 using a 100X objective. The effect of knocking down *bacA_CT_* on the production of infectious progeny was quantified by measuring the inclusion forming units (IFU) by assessing MOMP and GFP fluorescence **(F)**. The genomic DNA levels of uninduced and induced samples were also measured by qPCR to monitor the effect of *bacA*_CT_ knockdown on chlamydial replication **(G)**. Scale bar, 0.5 μm **(A, E)**; 2 μm **(B-D)**. **, p<0.05; ***, p<0.001; NS, Not significant.

To examine the degree to which knocking down *bacA_CT_* expression affected the chlamydial developmental cycle, we quantified the number of inclusion forming units (IFU), a proxy for EB generation, and the genomic DNA level during the developmental cycle. We quantified both the number of GFP-positive inclusions, indicative of bacteria carrying the CRISPRi plasmid encoding a constitutively expressed GFP, and those that labeled only with the antibody targeting MOMP, indicative of all bacteria. The ratio of GFP^+^ inclusions to MOMP^+^ inclusions gives a measure of plasmid stability, which can be negatively impacted by over-expressing, or potentially knocking down, a target gene (35). Importantly, no effect on chlamydial growth was detected after expressing the dCas9 alone **(Fig. S4B)**. Knocking down *bacA_CT_* transcripts resulted in roughly a two-fold decrease in MOMP^+^ IFUs and a three-fold decrease in GFP^+^ IFUs **(Fig. 4F)**. Furthermore, the genomic DNA levels were not significantly changed **(Fig. 4G)**, suggesting that *bacA*_CT_ knockdown does not grossly affect developmental cycle progression within the parameters of our experimental conditions. Overall, these results indicate that disrupting BacA_CT_ levels negatively impacts chlamydial morphology. These data support the hypothesis that BacA_CT_ is a morphological determinant in *Chlamydia*.

### BacA_CT_ knockdown can be complemented to restore normal morphology

We next sought to complement the knockdown phenotype by expressing either *bacA_CT__6xH* or *bacA_CT__mCherry* as a transcriptional fusion with *dCas9*. Under aTc-inducing conditions, the dCas9 will repress the chromosomal expression of *bacA_CT_* while the plasmid copy of *bacA_CT__6xH* or *bacA_CT__mCherry* is expressed. To measure transcriptional changes in chromosomal *bacA_CT_*, we introduced synonymous changes in the plasmid copy of *bacA_CT__6xH* or *bacA_CT__mCherry* that eliminated binding of the qPCR primers used as well as the gRNA binding site. When we performed this experiment, we measured a decrease in *bacA_CT_* transcripts as noted above **(Fig. 5A, B)**. Importantly, when we imaged bacteria from uninduced and induced conditions, we observed both dCas9 and BacA_CT_ labeling within RBs in the induced conditions only **(Fig. 5C, D)**. Critically, the RBs exhibited normal morphology under these conditions, thus indicating successful complementation of the knockdown phenotype. When taken together, our data demonstrate that BacA_CT_ 1) associates with the bacterial membrane, 2) fails to localize or interact with components of the bacterium’s cell division machinery, and 3) directly influences the size and shape of *Chlamydia’s* replicative form. Thus, we hypothesize that BacA_CT_ functions as a dynamic, membrane stabilizing agent during the pathogen’s growth and division process.

**Figure 5.**
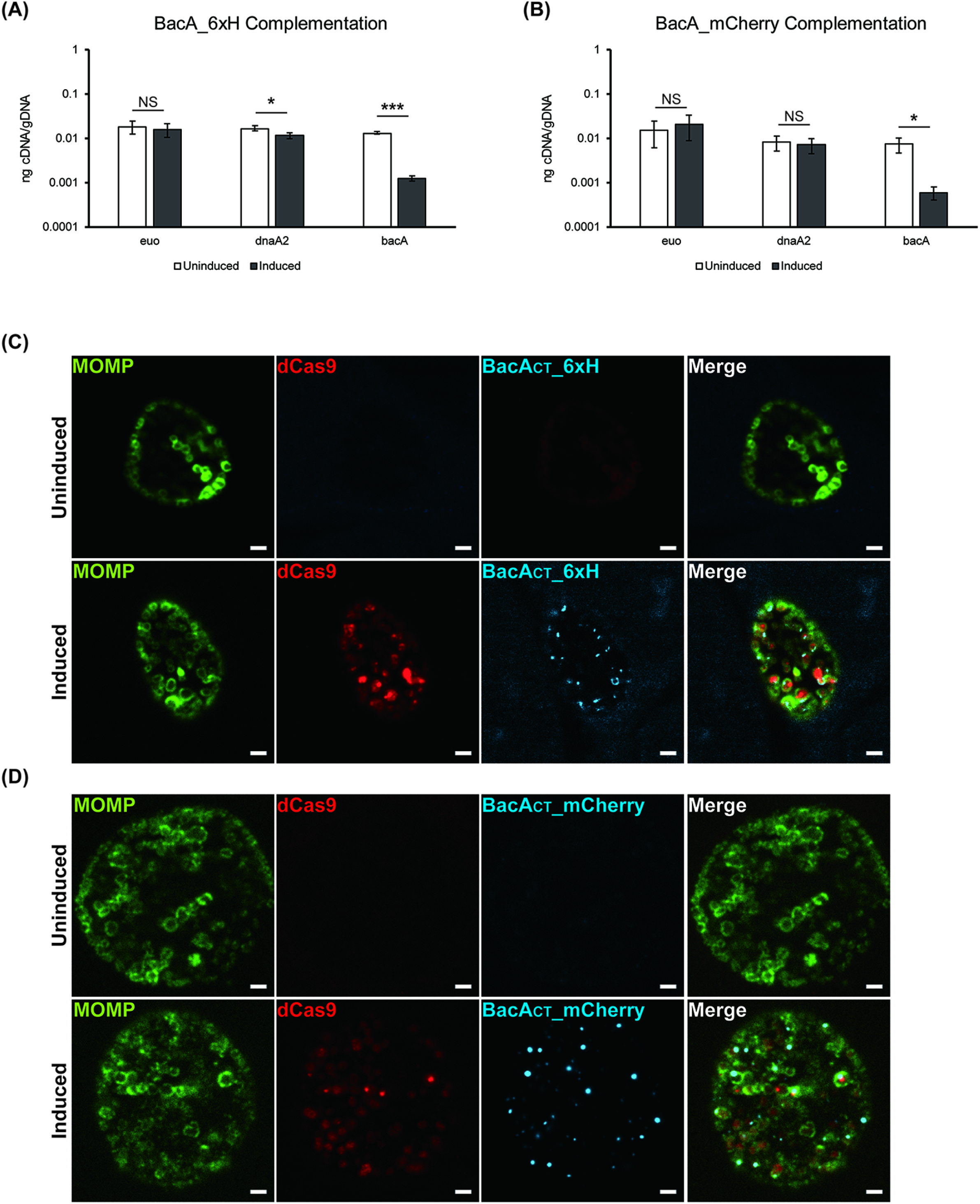
Ectopic expression of BacA_CT_ complements the morphological defects during knockdown of*bacA_CT_* expression in *C. trachomatis*. Constructs encoding an aTc-inducible dCas9 transcriptionally fused to BacA_CT__6xH or BacA_CT__mCherry and a constitutively expressed gRNA targeting *bacA* were made and transformed into *C. trachomatis* L2 (-pL2). Silent mutations were substituted into the qPCR primer binding and gRNA targeting sequence of *bacA_CT_* in the plasmid. Therefore, when RT-qPCR was performed, only endogenous *bacA_CT_* was detected. The transformants were infected into HeLa cells. The constructs were induced at 4 hpi, and the transcriptional level of endogenous *bacA_CT_* was measured by RT-qPCR at 24 hpi **(A, B)**. The RT-qPCR was performed as described in Materials and Methods. To confirm the expression of dCas9 and ectopic BacA_CT_, immunofluorescence assay was performed **(C, D)**. At 24 hpi, the samples were fixed with 1X DPBS containing 3.2% formaldehyde and 0.022% glutaraldehyde for 2 min. Afterwards, the samples were permeabilized with 90% methanol (MeOH). The staining of MOMP, dCas9, and BacA_CT__6xH was performed as described in Materials and Methods. Images were acquired on a Zeiss Imager.Z2 equipped with an Apotome2 using a 100X objective. Scale bar, 2 μm. *, p<0.05; ***, p<0.001; NS, Not significant

## Discussion

Bactofilins, or bacterial intermediate filament-like proteins, have been identified in many bacterial species and have been described to function in various cellular processes including maintenance of cell shape and morphology, establishment of cell polarity, and orientation of cell division proteins (11–14). Given the previously described functions of bactofilins in other bacteria, we initially hypothesized that BacA_CT_ would serve to direct the polarized division process of *Chlamydia*. However, based on our findings, our initial hypothesis of BacA_CT_ function is likely incorrect but does not preclude BacA_CT_ from functioning in the division process indirectly.

Although we failed to reveal a direct link between BacA_CT_ and cell division, we did observe that *Chlamydia* RBs exhibited distinct changes in their morphology when *bacA_CT_* expression was knocked down by CRISPR interference. Changes in morphology due to the loss of a bacterium’s bactofilin has been reported previously. The *P. mirabilis* protein, CcmA (curved cell morphology), was identified following the loss of curved cell morphology in a transposon mutant (36). Subsequently, other bactofilins were identified and found to be important in maintenance of bacterial cell morphology. For instance, loss of BacM in *M. xanthus* causes a kinked morphology (37), while the absence of CcmA in *H. pylori* results in curved rod morphology instead of the microbe’s distinct helical morphology (23), owing to the inability of PG remodeling proteins to properly localize. These bacteria are rod-shape species and have PG sacculi regulated in part by the function of their respective bactofilin. In contrast, members of the *Chlamydiaceae* do not have a PG sacculus, thus the exact mechanism by which BacA_CT_ regulates morphology is likely to be unique to these organisms. Further support for this comes from the fact that no homolog for BacA_CT_ is encoded in the genomes of ‘Chlamydia-related’ bacteria. These bacteria are predominantly endosymbionts and share *Chlamydia*’s biphasic developmental cycle. However, they physiologically differ from pathogenic *Chlamydia* species in that they retain a peptidoglycan sacculus (38).

Because pathogenic *Chlamydia* species lack sacculi, they must utilize an alternative means of maintaining cell shape and membrane integrity. In the microbe’s extracellular form, this is achieved via disulfide crosslinked, cysteine-rich outer membrane proteins. This structure is strong and rigid but does not provide for the membrane dynamics necessary for chlamydial cell growth and division. Peptidoglycan, and by extension its interacting partners in the divisome, and the Tol-Pal system (previously characterized in the Chlamydia-related organism *Waddlia chondrophila* (39)) have been shown to provide stability to the outer membrane in homologous systems, and BacA_CT_ may function similarly by stabilizing membrane regions outside the division plane, in conjunction with an as yet uncharacterized interacting partner in the periplasmic space **(Fig. 6)**. While this is one possibility, BacA_CT_’s function may prove inherently simpler. In addition to enabling bacterial cell division, peptidoglycan is essential for maintaining cell size and shape. Without a sacculus, the only way peptidoglycan can affect cell morphology in *Chlamydia* species is prior to division when its ring form places limitations on membrane expansion. However, upon septation, this structural scaffold is momentarily disassembled, leaving nothing to define the size and shape of a chlamydial RB in this intermediate state. BacA_CT_ rings could potentially bridge this critical stage in the division cycle and provide a degree of structural stability until a new peptidoglycan ring is formed. Although this study is the first to identify and characterize a bactofilin ortholog in *Chlamydia trachomatis*, there still remains much to uncover about the mechanism by which BacA_CT_ maintains cell morphology. By understanding the function of BacA_CT_ in *Chlamydia*, we will further understand the development of this unique obligate intracellular bacterium.

**Figure 6:**
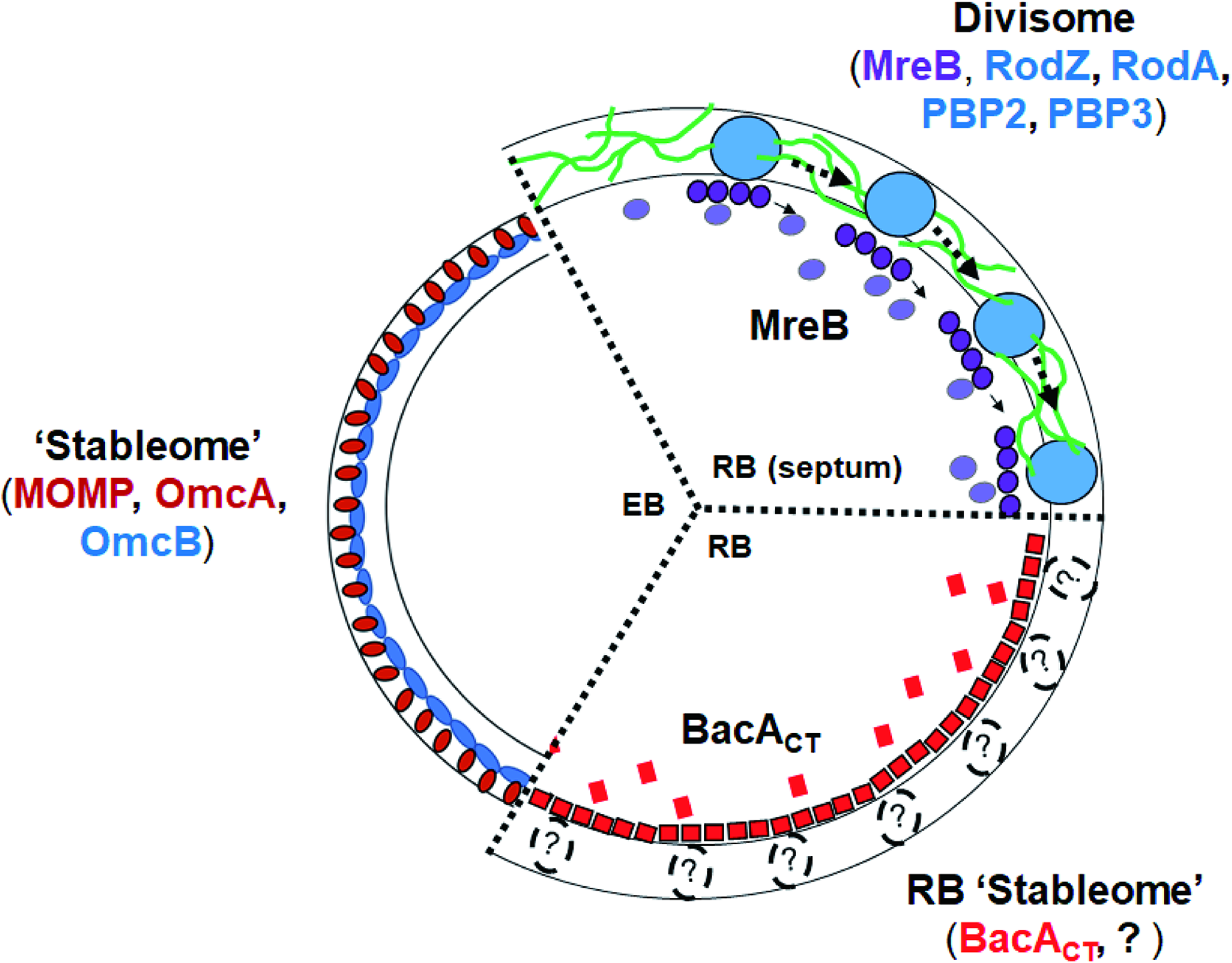
Membrane stabilization mechanisms are spatially restricted and differ between *Chlamydia’s* developmental forms. *Chlamydia* species rely on highly crosslinked outer membrane proteins (OmpB, OmpC) to maintain membrane integrity in their extracellular forms, however, this structure is rigid and does not allow for the membrane expansion and fluidity required during bacterial growth and division. Conversely, the microbe’s replicative form maintains peptidoglycan only at the septum, as this is the only place where the pathogen’s peptidoglycan synthases (divisome) localize. We hypothesize that BacA_CT_, likely in conjunction with other periplasmic elements, functions to stabilize the outer and inner membranes at areas other than the septum, effectively stabilizing specific areas of the periplasmic space and allowing transmembrane complexes to function in the absence of a sacculus.

## Materials and Methods

### Organisms and Cell Culture

McCoy (kind gift of Dr. Harlan Caldwell, NIH) and L2 mouse fibroblast cell lines and HeLa (ATCC, Manassas, VA) human epithelial cell line were cultured at 37°C with 5% CO_2_ in Dulbecco’s Modified Eagle Medium (DMEM; Invitrogen, Waltham, MA) containing 10% fetal bovine serum (FBS; Hyclone, Logan, UT) and 10 μg/mL gentamicin (Gibco, Waltham, MA). *Chlamydia trachomatis* serovar L2 (strain 434/Bu) lacking the endogenous plasmid (-pL2; kind gift of Dr. Ian Clarke, Univ. Southampton) was infected and propagated in McCoy cells for use in transformations. HeLa cells were infected with chlamydial transformants in DMEM containing 10% FBS, 10 μg/mL gentamicin, 1 U/mL penicillin G, and 1 μg/mL cycloheximide. All cell cultures and chlamydial stocks were routinely tested for *Mycoplasma* contamination using the Mycoplasma PCR detection kit (Sigma, St. Louis, MO). *Chlamydia trachomatis* strain L2/434 Bu was obtained from the laboratory of Dr. Anthony Maurelli (University of Florida). *E. coli* TOP10 cells were used for subcloning and propagating of the shuttle vector, pREF100, whereas *E. coli* NEB10β (New England Biolabs, Ipswich, MA) cells were used for pBOMB and pTLR2 vectors. BACTH vectors were cloned and propagated in *E. coli* NEB5αI^q^ cells (NEB). *E. coli* was grown at 30°C or 37°C in LB broth. Chlamydial transformations and EB recovery assays were performed in L2 or McCoy cells (mouse fibroblast-derived cell lines) while imaging infection studies were performed in HeLa cells. All chemicals and antibiotics were obtained from Sigma unless otherwise noted.

### Bioinformatics Analysis

Sequences for *Chlamydia trachomatis* serovar L2/434 and other chlamydial species, *Caulobacter crescentus, Myxococcus xanthus,* and *Thermus thermophilus* were obtained from the NCBI database (https://www.ncbi.nlm.nih.gov/)(40). Protein sequence alignment was performed using Clustal Omega website (https://www.ebi.ac.uk/Tools/msa/clustalo/)(41) and the ESPript3 program (http://espript.ibcp.fr)(42). The 3D-structure prediction was used by Swiss modeling program (https://swissmodel.expasy.org/) (43) and RaptorX web-based applications. Visualization of pathogenic chlamydial BacA structural predictions was performed with the UCSF Chimera imaging, analysis, and visualization suite (44).

### Cloning

The plasmids and primers used in this study are described in Supplemental Table 1. The wild-type *Chlamydia trachomatis ct276* gene was amplified by PCR with Phusion DNA polymerase (NEB) using 100 ng *C. trachomatis* L2 genomic DNA as a template. Some gene segments were directly synthesized as a gBlock fragment (Integrated DNA Technologies, Coralville, IA). The PCR products were purified using a PCR purification kit (Qiagen, Hilden, Germany). The HiFi Assembly reaction master mix (NEB) was used according to the manufacturer’s instructions in conjunction with plasmids pASK-GFP-mKate2-L2 (kind gift of Dr. P. Scott Hefty, Univ. Kansas) cut with FastDigest AgeI and EagI (Thermofisher, Waltham, MA), pBOMB4-Tet (kind gift of Dr. Ted Hackstadt) cut with EagI and KpnI, or pKT25 or pUT18C cut with BamHI and EcoRI, depending on the construct being prepared. The CRISPRi plasmid targeting *ct276* was constructed by amplifying the *S. aureus* dCas9 from pX603-AAv-CMV::NLS-dSaCas9(D10A,N580A)-NLS-3xHA-bGHpA (a kind gift of Dr. Feng Zhang; Addgene plasmid 61594 (32)) and inserting it into a modified pBOMB4-Tet vector (Ouellette et al., *in prep*). A gRNA specific to the 5’ end of *ct276* was then introduced or not into the BamHI site of the Sa_dCas9 encoding plasmid (see Suppl. Table S1). All plasmids were also dephosphorylated with FastAP (ThermoFsher). The products of the HiFi reaction were transformed into either NEB10β (for chlamydial transformation plasmids) or NEB5αI^q^ (for BACTH) competent cells (NEB) and plated on appropriate antibiotics, and plasmids were subsequently isolated from individual colonies grown overnight in LB broth by using a mini-prep kit (Qiagen). For chlamydial transformation, the constructs were transformed into *dam*^-^ *dcm*^-^ competent cells (NEB) and purified as de-methylated constructs. To create the mCherry fusion construct, mCherry was amplified from pBOMB-tet-mCherry (45) and inserted into the expression vector pREF100 (19) (a kind gift from Dr. Anthony Maurelli). The *bacA_CT_* and *mCherry* amplicons were joined together by overlapping PCR. The insert PCR product and pREF100 were double digested with NotI and PstI (NEB). Digested PCR products were ligated into the pREF100 shuttle vector downstream of an anhydrous tetracycline (aTc) inducible promoter. pREF100:bacA-His or pREF100:bacA-mCherry plasmids were transformed into chemically competent TOP10 *E. coli*. Plasmids were extracted, sequenced, and transformed into either *C. trachomatis* strain L2/434 Bu or strain −pL2 via exposure to CaCl_2_, described below.

### Transformation of *Chlamydia trachomatis*

McCoy or HeLa cells were plated in a six-well plate the day before beginning the transformation procedure. *Chlamydia trachomatis* serovar L2 without plasmid (-pL2) (kind gift of Dr. Ian Clarke) was incubated with 2 μg demethylated plasmid in Tris-CaCl_2_ buffer (10 mM Tris-Cl, 50 mM CaCl_2_) for 30 min at room temperature. During this step, the McCoy / HeLa cells were washed with 2 mL Hank’s Balanced Salt Solution (HBSS) media containing Ca^2+^ and Mg^2+^ (Gibco). After that, cells were infected with the transformants in 2 mL HBSS per well. The plate was centrifuged at 400 *x g* for 15 min at room temperature and incubated at 37°C for 15 min. The inoculum was aspirated, and DMEM containing 10% FBS and 10 μg/mL gentamicin was added per well. At 8 h post infection (hpi), the media was changed to media containing 1 μg/mL cycloheximide and either 1 or 2 U/mL penicillin G (for pBOMB4-Tet constructs) or 100 μM spectinomycin (for pREF100 constructs), and the plate was incubated at 37°C until 48 hpi. At 48 hpi, the transformants were harvested and used to infect a new McCoy or HeLa cell monolayer. These harvest and infection steps were repeated every 44-48 hpi until mature inclusions were observed. Chlamydial transformants were then plaque purified at least once (to ensure clonal populations) and propagated to high titer stocks for use in subsequent experiments.

#### Viable EB / IFU Assay

L2 mouse fibroblast cells were infected with *C. trachomatis* L2 434/Bu: pREF100: bacA_CT_-mCherry or bacA_CT_-His at a MOI of ~1 in a 24 well plate with 0, 1, or 10 ng/ml of aTc for 44 hours. EBs were then mechanically harvested with glass beads and 200 μl of 1X sucrose-phosphate-glutamate (SPG) buffer. BacA_CT_-mCherry EBs were stored at −80°C, while BacA_CT_-His EBs were used immediately to re-infect a new monolayer of cells. To re-infect cells, EBs were serially diluted and placed on L2 cell monolayers, seeded in a 96 well plate. At 24 hpi, 20 fields of view of GFP^+^ inclusion forming units (IFU) were counted in duplicate for each treatment condition. Three biological replicates were performed.

#### Labeling of chlamydial peptidoglycan and MreB

HeLa cells were infected with *C. trachomatis* L2 434/Bu: pREF100:bacA_CT_-His or *C. trachomatis* L2 434/Bu: pREF100:bacA_CT_-mCherry and induced with 10 ng/ml of aTc at 2hpi. At 10.5, 21, and 44 hpi, cells were washed 3 times with 1X PBS, permeabilized with methanol and 0.5% Triton X (5 mins each) prior to blocking with 3% Bovine Serum Albumin (BSA) for 1 hour and antibody labeling. Peptidoglycan was metabolically labelled with D-amino acid dipeptide probes (1mM EDA-DA) and click chemistry as previously described (9, 10). His-tagged BacA_CT_ was labeled with anti-His primary antibody (mouse) and secondary antibody anti-mouse AlexFluor 594), at the dilutions 1:200 and 1:2000, respectively. Similarly, to label MreB and BacA, HeLa cells were infected with *C. trachomatis* L2 434/Bu: pREF100:bacA_CT_-mCherry and induced with 10 ng/ml of aTc. At 21 hpi, cells were washed 3 times with 1X PBS, permeabilized with methanol and 0.5% Triton X (5 mins each) prior to blocking with 3% Bovine Serum Albumin (BSA) for 1 hour and antibody labeling. Major Outer Membrane Protein (MOMP) was labeled with anti-MOMP primary antibody (goat) and secondary antibody (donkey anti-goat AlexaFluor blue) at the dilutions 1:500 and 1:1000 respectively. MreB was labeled with rabbit, anti-MreB primary antibody (30) (generously provided to us by Dr. Scott Hefty of Kansas University) and anti-rabbit secondary conjugated to AlexaFluor 488.

#### Live imaging confocal microscopy studies

HeLa cells seeded in chamber wells were infected C. *trachomatis* L2 434/Bu: pREF100:bacA_CT_-mCherry and induced with 10ng/ml of aTc at 2 hpi. Cells were imaged utilizing a Zeiss 710 confocal microscope under tissue culture conditions (see above). zStacks were acquired at a variety of time points post infection, and were used to illustrate changes occurring using single imaging planes or to generate maximum intensity projections. All video editing was done using Zen Black software (Zeiss).

### Indirect Immunofluorescence (IFA) Microscopy

HeLa cells were seeded in 24-well plates on coverslips at a density of 10^5^ cells per well the day before infection. Chlamydial strains expressing wild-type or truncated Ct276 with a six-histidine tag at the C-terminus were used to infect HeLa cells in DMEM media containing penicillin G and cycloheximide. At 6 hpi, 10 nM anhydrotetracycline (aTc) was added. At 10.5 hpi, the coverslips of infected cells were washed with DPBS and fixed with fixing solution (3.2% formaldehyde and 0.022% glutaraldehyde in DPBS) for 2 min. The samples were then washed three times with DPBS and permeabilized with ice-cold 90% methanol for 1 min. Afterwards, the fixed cells were labeled with primary antibodies including goat anti-major outer-membrane protein (MOMP; Meridian, Memphis, TN) and mouse or rabbit anti-six histidine tag (Genscript, Nanjing, China and Abcam, Cambridge, UK, respectively). For experiments using CRISPR interference, a rabbit anti-Sa_Cas9 was used (Abcam). Secondary antibodies donkey anti-goat antibody (594), donkey anti-rabbit antibody (647), and donkey anti-mouse (488) were used to visualize the primary antibodies. The secondary antibodies were obtained from Invitrogen. Coverslips were observed by using either a Zeiss LSM 800 confocal microscope or Zeiss Apotome.2.

### BACTH Assay

The pKT25 and pUT18C vectors encoding the genes of interest or empty vectors were co-transformed into competent DHT1 *E. coli*, an adenylate cyclase-deficient strain, and spread onto M63 minimal media plates containing 50 μg/mL ampicillin, 25 μg/mL kanamycin, 0.5 mM IPTG, 0.2% maltose, 40 μg/mL X-gal, and 0.04% casamino acids. The plates were incubated at 30°C for up to seven days. Blue colonies indicate positive interactions whereas no growth or small white colonies indicate no interactions. β-galactosidase assays were performed as described elsewhere on overnight cultures grown in M63 minimal medium broth containing the same supplements except for X-gal (17, 29).

### Transcriptional Analyses in *Chlamydia*

HeLa cells were infected with the *C. trachomatis* L2 transformants carrying the Sa_dCas9 +/-gRNA plasmid. At 4hpi, Sa_dCas9 expression was induced. Total RNA and DNA were collected from replicate wells using Trizol (Invitrogen) and DNeasy Tissue (Qiagen) kit as described elsewhere (18). After DNasing total RNA, cDNA was synthesized using Superscript III reverse transcriptase (Invitrogen), and equal volumes of cDNA were used in qPCR reactions (18). Similarly, equal mass of DNA was used from each sample in qPCR reactions to quantify chlamydial genomes, which were used to normalize transcript data (46).

## ACKNOWLEDGEMENTS

The authors would like to thank the following individuals for providing reagents used in this study: Dr. Harlan Caldwell (National Institute of Health), Dr. Ian Clarke (University of Southampton), Dr. Ted Hackstadt (Rocky Mountain Labs/NIH), Dr. Anthony Maurelli (University of Florida), Dr. Michael VanNieuwenhze (University of Indiana), Dr. Scott Hefty (University of Kansas), and Dr. D. Scott Merrell (Uniformed Services University). This work was supported by grants from the National Institute for General Medical Science (R35GM124798-01 to SPO and R35GM138202-01 to GWL) within the National Institutes of Health as well as a faculty start-up package awarded by the Uniformed Services University to GWL. The funders had no role in study design, data collection and interpretation, or the decision to submit the work for publication. The University of Nebraska Medical Center Advanced Microscopy Core Facility receives partial support from the National Institute for General Medical Science INBRE −P20 GM103427 and COBRE −P30 GM106397 grants, as well as support from the National Cancer Institute (NCI) for The Fred & Pamela Buffett Cancer Center Support Grant-P30 CA036727 and the Nebraska Research Initiative.

The opinions and assertions expressed herein are those of the author(s) and do not necessarily reflect the official policy or position of the Uniformed Services University or the Department of Defense.

## AUTHOR CONTRIBUTIONS

MRB, JL, GWL, and SPO designed and implemented experiments and wrote the manuscript. MRB, JL, JVC, GWL, and SPO interpreted data. MRB, JL, and GWL prepared figures. JVC, GWL, and SPO edited the manuscript. JVC, GWL, and SPO provided oversight. GWL and SPO secured funding for the work.

## Supplemental Figure Legends

**Table S1: List of primers, plasmids, and strains used in the study**.

**Figure S1: BacA_CT_ contains a β-helical core domain that is universally conserved in all bactofilins. (A)** Amino acid sequence alignment of pathogenic *Chlamydia sp.* BacA. β-helical core domain is delineated by the blue bar. **(B)** Structural predictions of BacA_CT_ encoded pathogenic *Chlamydia* species.

**Figure S2. Induction of BacA_CT_ fusion constructs does not affect chlamydial development.** L2 mouse fibroblast cells were infected with (A) *C. trachomatis* L2 434/Bu: pREF100:BacA_CT_-mCherry or (B) BacA_CT_-His at an MOI of 1 in a 24 well plate with 0, 1, or 10 ng/ml of aTc for 44 hours. EBs were then mechanically harvested, serially diluted, and used to infect new L2 cell monolayers in a 96 well plate. At 24 hpi 20 fields of view of GFP^+^ inclusion forming units (IFU) were counted in duplicate for each treatment condition. The solid line represents the mean of three biological replicates and the dotted line represents the assay’s limit of detection.

**Figure S3. BacA_CT_ ring morphology observed in fixed and living cells via super resolution microscopy. (A)** Cells were grown in the presence or absence of 10 ng/mL anhydrotetracycline. At 18 hours post infection, cells were fixed, permeabilized, and labeled with polyclonal antibodies against mCherry (red) and *Chlamydia’s* Major Outer Membrane Protein, MOMP (blue). Representative Z-Stack images were taken via super-resolution microscopy. **(B)** HeLa cells were infected with *C. trachomatis* L2 434/Bu: pREF100:BacA_CT_-mCherry with 10 ng/ml aTc induction. GFP^+^ *Chlamydia* (green) and BacA_CT_-mCherry (red). Images were taken at 37 hpi. (**C**) MreB does not co-localize with BacA_CT_. Super resolution microscopy of *C. trachomatis* expressing BacA_CT_-mCherry (red) 18 hours post infection. Cells were fixed, permeabilized, and immunolabeled utilizing polyclonal antibodies to mCherry and the chlamydial cell division protein, MreB. Images are max intensity projections and representative of ~20 imaged fields. Scale bar, 1 μm.

**Figure S4. The effect of dCas9 over-expression without gRNA in *C. trachomatis.* (A)***C. trachomatis* serovar L2 transformed with a vector encoding dCas9 only (i.e. without a gRNA) was used to infect HeLa cells. At 4 hpi, dCas9 was induced with 20 nM aTc, and the transcriptional levels of *euo*, *dnaA2*, and *bacA* were measured by qPCR at 24 hpi **(A).** Also, to check the expression of dCas9, the immunofluorescence assay was performed at 24 hpi **(B)**. The staining was performed as described in Materials and Methods. Images were acquired on a Zeiss Imager.Z2 equipped with an Apotome2 using a 100X objective. Scale bar, 2 μm. NS, Not significant

**Figure S5. Knockdown of BacA_CT_ does not alter peptidoglycan ring formation.** Images are separated channels from data shown in main Figure 4E.

**Supp. Video S1. Time lapse microscopy of BacA_CT_ in living cells.** HeLa cells were infected with *C. trachomatis* L2 434/Bu: pREF100:BacA_CT_-mCherry with 10 ng/ ml aTc induction. GFP+ *Chlamydia* (green) and BacA_CT_-mCherry (red). Images were taken at 40 hpi and inclusion is approximately midway through the microbe’s developmental cycle as numerous RBs are still visible. Images represent a cross-section of infected cells and a single imaging plane.

**Supp. Video S2. Time lapse microscopy of BacA_CT_ in living cells.** HeLa cells were infected with *C. trachomatis* L2 434/Bu: pREF100:BacA_CT_-mCherry with 10 ng/ml aTc induction. GFP+ *Chlamydia* (green) and BacA_CT_-mCherry (red). Images were taken at 40 hpi and inclusion is approximately at the end of the microbe’s developmental cycle. The few BacA_CT_+ RBs that remain exhibit a polar orientation within the inclusion, while the vast majority of the inclusion space is filled with rapidly-moving, GFP+ BacA_CT_-EBs. Images represent a cross-section of infected cells and a single imaging plane.

**Supp. Video S3. Time lapse microscopy of BacA_CT_ in living cells.** HeLa cells were infected with *C. trachomatis* L2 434/Bu: pREF100:BacA_CT_-mCherry with 10 ng/ml aTc induction. GFP+ *Chlamydia* (green) and BacA_CT_-mCherry (red). Images were taken at approximately 40 hpi. Images represent a cross-section of infected cells and a single imaging plane.

**Video S4. Time lapse microscopy of BacA_CT_ transitioning from puncta to ring-like form.** HeLa cells were infected with *C. trachomatis* L2 434/Bu: pREF100:BacA_CT_-mCherry with 10 ng/ml aTc induction. GFP+ *Chlamydia* (green) and BacA_CT_-mCherry (red). Each imaging frame represents a 3D maximum intensity projection compiled from 15 individual imaging fields.

